# Wnt activation prevents epileptogenic hippocampal remodeling in animal models of unilateral and bilateral temporal lobe epilepsy

**DOI:** 10.64898/2026.05.05.722655

**Authors:** Cora Helton, Nicole Rodgers, Kunal Gupta

**Author notes:** Corresponding author: Kunal Gupta MD PhD. Declarations of interest: The authors declare no competing interests.

## Abstract

Temporal lobe epilepsy (TLE) is a heterogeneous disorder with most clinical presentations involving unilateral or bilateral hippocampal seizure onsets. Antiseizure medications are often ineffective for TLE, and epilepsy surgery can have variable outcomes. Risk factors for TLE are readily identifiable and typically precede chronic epilepsy, providing a window of opportunity for preventative treatments. However, there are currently no clinically approved anti-epileptogenic therapies. In this study, we investigate the role of Wnt signaling in epileptogenesis using two mouse TLE models, the intrahippocampal kainate model of unilateral TLE (IHK), and the intraperitoneal kainate model of bilateral TLE (IPK). We specifically examined adult-born immature dentate granule cells as these cells have been heavily implicated in the pathogenesis of TLE and clinical TLE is typically initiated in adulthood. We observed that adult-born immature dentate granule cells undergo pathological morphological changes during epileptogenesis in both the IHK and IPK models of TLE. When compared across epileptogenic zones, however, these changes differed between the two models. Wnt signaling also decreased in these cells in epileptic mice during the epileptogenic period. When mice were treated with SB415286, a highly selective Wnt activator, Wnt signaling in immature dentate granule cells was restored to baseline levels and pathological remodeling changes were reduced in both models. These data therefore suggest that a reduction in Wnt signaling in immature dentate granule cells plays an etiological role in epileptogenesis, and that restoring Wnt signaling using Wnt activating drugs or alternative agents may have therapeutic potential as an anti-epileptogenic strategy in TLE.

## Introduction

Temporal lobe epilepsy is the most common form of acquired focal epilepsy in adults, comprising approximately two-thirds of focal epilepsies ^1^. Of these, mesial or hippocampal onset temporal lobe epilepsy (MTLE) comprises 80% of temporal lobe epilepsies overall ^2^ and is characterized by a range of clinical presentations, including unilateral and bilateral hippocampal seizure onsets. TLE is typically managed with anti-seizure medications, however, many are poorly tolerated and worsen functional impairment ^3,4^. Furthermore, MTLE is medically refractory in up to one-third of patients ^5^, with higher rates of drug resistance in the setting of hippocampal sclerosis, the most common pathological finding in MTLE ^1,6^. Patients with drug resistant MTLE may be candidates for epilepsy surgery, however, surgical treatments and outcomes for unilateral and bilateral TLE differ greatly. There are several resective and ablative surgical options for unilateral TLE with comparatively higher rates of seizure freedom ^7,8^. The surgical treatment of bilateral TLE, however, is considerably more limited due to cognitive deficits associated with bilateral hippocampal resection ^9,10^ and is therefore typically limited to neuromodulation, which has comparatively lower rates of seizure freedom ^11,12^. There is therefore a vital need for improved management options for patients with different presentations of mesial temporal lobe epilepsy.

Epilepsy is acquired by a process called epileptogenesis; in this process, neurons are remodeled to generate epileptic circuits. Several high-risk events are known to induce epileptogenesis ^13,14^, and it is well established that after a single seizure, the risk of a recurrent seizure is as high as 43% after 2-years ^13–15^. A targeted intervention during this latent period between first seizure and seizure recurrence may be able to delay or prevent the onset of chronic epilepsy, however, there are currently no clinical therapies available that can prevent epileptogenesis. Notably, risk factors for unilateral and bilateral MTLE are shared, including structural abnormalities, infection, inflammation, stroke, traumatic brain injury and others ^16–20^, suggesting that these forms of MTLE may share common underlying etiopathogenesis and molecular signaling. Therefore, anti-epileptogenic interventions for MTLE may need to provide preventative efficacy for both unilateral and bilateral forms of TLE, as risk factors alone may be insufficient to determine which form of MTLE may ensue.

Rodent epilepsy models provide valuable opportunities to investigate the mechanisms responsible for epileptogenesis and test novel anti-epileptogenic therapies for clinical translation. Kainic acid is a well-established chemoconvulsant that can be administered either focally or systemically to induce epileptogenesis ^21^. The intrahippocampal kainate (IHK) model utilizes the direct injection of kainic acid into the hippocampus to induce unilateral hippocampal sclerosis and unilateral TLE. In the intraperitoneal kainate model (IPK), kainic acid is administered systemically by intraperitoneal injection, resulting in bilateral hippocampal remodeling and bilateral TLE. Notably, previous IPK models that reported low rates of survival or neurodegeneration utilized single injections of high dose kainate ^22–24^; newer repeated low dose models used in this study reliably induce acute status epilepticus followed by epileptogenesis and chronic epilepsy ^25,26^. In both models, mice reliably develop acute behavioral and electrographic status epilepticus followed by epileptogenesis within 10-14 days ^26–29^.

There is growing evidence that changes in Wnt signaling occur in the hippocampus during epileptogenesis ^30–33^. The Wnt pathway is comprised of 19 secreted Wnt ligands, which bind a family of frizzled receptors and co-receptors and signal to canonical beta-catenin dependent and independent downstream pathways ^34^. Beta-catenin, the principal effector of canonical Wnt pathway, is negatively regulated by GSK3b, which itself can be inhibited to activate canonical Wnt signaling ^35,36^. Epileptogenesis in TLE is characterized by hippocampal dentate granule cell remodeling, which alters their connectivity and excitability resulting in the formation of epileptic networks. These processes are regulated by Wnt signaling and, in our previous work, we have shown that Wnt signaling is downregulated during epileptogenesis ^32^. Furthermore, Wnt activation using Chir99021 during epileptogenesis in the IHK model of unilateral MTLE reduces seizure burden and prevents pathological granule cell dendrite remodeling associated with epileptogenesis ^27^. Notably, however, Chir99021 treatment did not alter other MTLE-associated histological markers, including granule cell dispersion, migration or neurogenesis, thereby retaining several pathological remodeling features consistent with TLE ^37–39^. Moreover, the effect of Wnt activation was only tested in the IHK model of unilateral TLE.

In this study, we investigate the effect of a highly selective Wnt activating compound SB415286 on hippocampal epileptogenic remodeling in both the IHK model of unilateral TLE and the IPK model of bilateral TLE. SB415286 is a brain-penetrant Wnt activator from the maleimide class of GSK3b inhibitors with minimal off-target kinase modulating effects ^36,40^. Our data demonstrate that Wnt activator SB415286 prevents epileptogenic remodeling of hippocampal dentate granule cells in both unilateral and bilateral MTLE models, with greater preventative effects than Chir99021 after IHK. We also assessed Wnt activation in adult-born immature granule cells and demonstrate that Wnt signaling decreases in these dentate granule cells during epileptogenesis and that Wnt activator SB415286 restores Wnt signaling activity to baseline levels in these cells. Notably, these neurons are specifically implicated in epileptogenesis in TLE ^27,41–43^. Our findings therefore provide evidence that Wnt activation may provide an anti-epileptogenic therapeutic approach for two distinct clinical presentations of TLE and may act mechanistically by altering Wnt signaling in adult-born immature dentate granule cells.

## Methods

### Animal husbandry

Animal experiments were performed using 8–10-week-old wild-type C57Bl6/J mice (Jax #000664) and C57BL/6J-Tg(POMC-eGFP)1Low/J transgenic mice (JAX #009593). Mice were housed in group cages, with access to food and water *ad libitum*. In the POMC-GFP line, adult-born immature dentate granule cell neurons express enhanced green-fluorescent protein (GFP) for 2-weeks after birth with over 90% overlap in labeling with doublecortin (DCX) ^41,44^. These studies focused on immature dentate granule cells due to their contribution to temporal lobe epileptogenesis ^42^. Mice were housed in group cages at constant temperature and humidity in a 14h light/10h dark cycle with food and water *ad libitum*. In all experiments, male and female mice were used 1:1; data represent pooled results unless specified. All animal procedures were performed in accordance with the NIH *Guide for the care and use of laboratory animals* and were approved by the Institutional Animal Care and Use Committee (AUA #8273).

### Intrahippocampal kainate model of unilateral temporal lobe epilepsy (IHK)

Mice underwent unilateral intrahippocampal injection with kainate (Tocris, cat no. 7065) to induce unilateral mesial temporal lobe epilepsy ^27,32^; intrahippocampal saline injection was used for control animals. Mice were induced and maintained under anesthesia with inhaled isoflurane. The head was secured in stereotactic apparatus (Stoelting), the scalp was shaved and sterilized. A midline incision was made and stereotactic coordinates obtained relative to bregma. 100nl of saline or kainate (2mg/ml, 0.2µg / 1nmol in 100nl) (Tocris, cat no. 7065) were injected into the CA3 region of the hippocampus (X 1.8, Y -2.1, Z -1.9), the needle was retained for 2 minutes prior to stepped removal to prevent reflux. The skin was closed using 4-0 prolene sutures, and mice were recovered in a warmed chamber. No mice that received intrahippocampal saline control injection met criteria for status epilepticus.

### Intraperitoneal kainate model of bilateral temporal lobe epilepsy (IPK)

Bilateral temporal lobe epilepsy was induced by repeated low dose injection with intraperitoneal kainate; equivalent injections with intraperitoneal saline were used for control animals. Mice underwent repeated low-dose intraperitoneal kainate injection until the induction of convulsive status epilepticus; this model reliably induces epileptogenesis ^25,26,45^. Mice were loaded with 10mg/kg IP kainate (Tocris, cat no. 7065) and then re-dosed with 5mg/kg IP kainate every 30 minutes until they entered convulsive status epilepticus. No further kainate injections were administered after mice met criteria for status epilepticus. No mice that received intraperitoneal saline control injections met criteria for status epilepticus.

### Seizure scoring

Seizures were scored behaviorally for 2-hours after completion of surgery in the IHK model and after initiation of injections in the IPK model by modified Racine scale; stages 1 and 2 demonstrated freezing, mastication and head nodding, stage 3 demonstrated unilateral forelimb clonus, stage 4 demonstrated bilateral forelimb clonus and rearing, stage 5 demonstrated rearing and falling, and stage 6 demonstrated “popcorn” type seizures ^46^. No saline injected mice exhibited seizures during the study. Mice were included in the epilepsy group if they demonstrated convulsive status epilepticus, defined as continuous Racine 3-6 seizures for at least 30 minutes without recovery, during the observation period. These criteria meet guidelines for severe SE, which reliably causes epileptogenesis in both the IHK model ^27,29^ and in the repeated low dose IPK model ^25^.

### Drug treatments

Canonical Wnt activator SB415286 (Selleck Chemicals, cat no. S2729) was dissolved in 10% DMSO / 45% polyethylene glycol 400 / 45% normal saline. The same solvent without SB415286 was used for vehicle controls. SB415286 was dosed at 10mg/kg/day IP ^47–49^. SB415286 and vehicle treatments were dosed by weight for an IP injection volume of 100-150µl based on an average adult mouse weight of 20-30g. Mice were dosed daily, once every 24 hours, until euthanasia.

### Histology and immunohistochemistry

Mice were euthanized and transcardially perfused with PBS followed by 4% paraformaldehyde for fixation. Brains were extracted, post-fixed for 24 hours in 4% PFA and dehydrated in 30% sucrose. Coronal brain slices were obtained by cryosection (50µm), random slices are blocked for 1 hour with 5% goat serum/0.5% Triton-X/PBS and immunolabeled with primary antibodies (guinea-pig anti-doublecortin 1:250, mouse anti-active beta catenin 1:400, rabbit anti-Axin2 1:200) and fluorophore-conjugated secondary antibodies (AlexaFluor, 1:1000) in 2% goat serum/0.5% Triton-X/PBS, each applied for 24hrs at 4°C. Nuclei were counterstained with DAPI (1:10,000) prior to mounting (Vectashield Hardset). Images were obtained by confocal microscopy (20µm stacks, 1.0µm steps, Axin2: 63x, POMC-GFP: 20x).

### Histological quantification

Granule cell layer width, granule cell count, migration and primary dendrite angle were quantified in individual confocal image slices across full stacks using blinded image sets (ImageJ, NIH). A POMC-GFP+ cell was considered within the granule cell layer if the center of the soma was located within 10µm of the inferior border of the granule cell layer to include the subgranular zone ^41^. Cell counts were normalized to granule cell layer volume. Migration was calculated from the inferior border of the granule cell layer to the midpoint of the granule cell body. For primary dendrite angle, the orientation of the primary dendrite for each POMC-GFP+ neuron was measured with respect to the borders of the granule cell layer perpendicular to each individual cell. The apex of the dentate gyrus was designated as 0° and the hilus designated as 180° ^27,43^. Example traces are provided in Supplemental Figures 1 and 2. Axin2 fluorescence intensity was quantified in ImageJ by measurement of mean intensity, selecting cell somata as regions of interest for quantification; data were normalized for background secondary antibody staining in the inner molecular layer for each image to control for variations in background fluorescence intensity across individual animals. Axin2 is a well-established quantifiable marker of Wnt activity ^50–52^. For the IHK model, data are reported ipsilateral and contralateral to the side of intrahippocampal injection. For the IPK model, both hemispheres were quantified and averaged as no statistically significant differences were identified between hemispheres in this model (Supplemental Figure 3).

### Statistics

Statistical analyses were performed using Prism 10 (GraphPad, La Jolla CA). Data are presented as mean ± standard error. For each experiment, *n* per group was 6, male and female mice were used equally, data were pooled for both sexes. Comparisons between two groups were performed with paired (within animal) and unpaired (between animals) Student’s t-test. Comparisons of three or more groups were performed by analysis of variance analysis (ANOVA) with Tukey’s multiple comparisons test. Primary dendrite angle distributions were analyzed by Kolmogorov-Smirnov test. All tests were two-sided, *p*<0.05 was considered statistically significant. Significance thresholds are displayed as \**p*<0.05, \*\**p*<0.01, \*\*\**p*<0.001, \*\*\*\**p*<0.0001.

## Results

We quantified canonical Wnt activation in the hippocampal dentate gyrus after SB415286 treatment. SB415286 is a highly selective CNS-penetrant anilinomaleimide GSK3b inhibitor with established beta-catenin activating properties ^36^ and a half-life of 20 hours ^47^. Mice were administered a single intraperitoneal dose of SB415286, and canonical Wnt pathway activity was quantified in hippocampal dentate granule cells 3-hours and 24-hours after treatment (Figure 1A). Wnt activity in the granule cell layer was identified by immunoreactivity for Axin2, a beta-catenin activity dependent gene ^50,52^ (Figure 1B). Wnt activity increased 5±0.94 fold 3-hours after SB415286 treatment and remained 7±1.25 fold increased after 24-hours, compared to vehicle treated controls (Figure 1C, *p*<0.01). These data demonstrate that systemic treatment with SB415286 induces Wnt activation in dentate granule cells, which was sustained at least 24-hours after a single dose.

**Figure 1.**
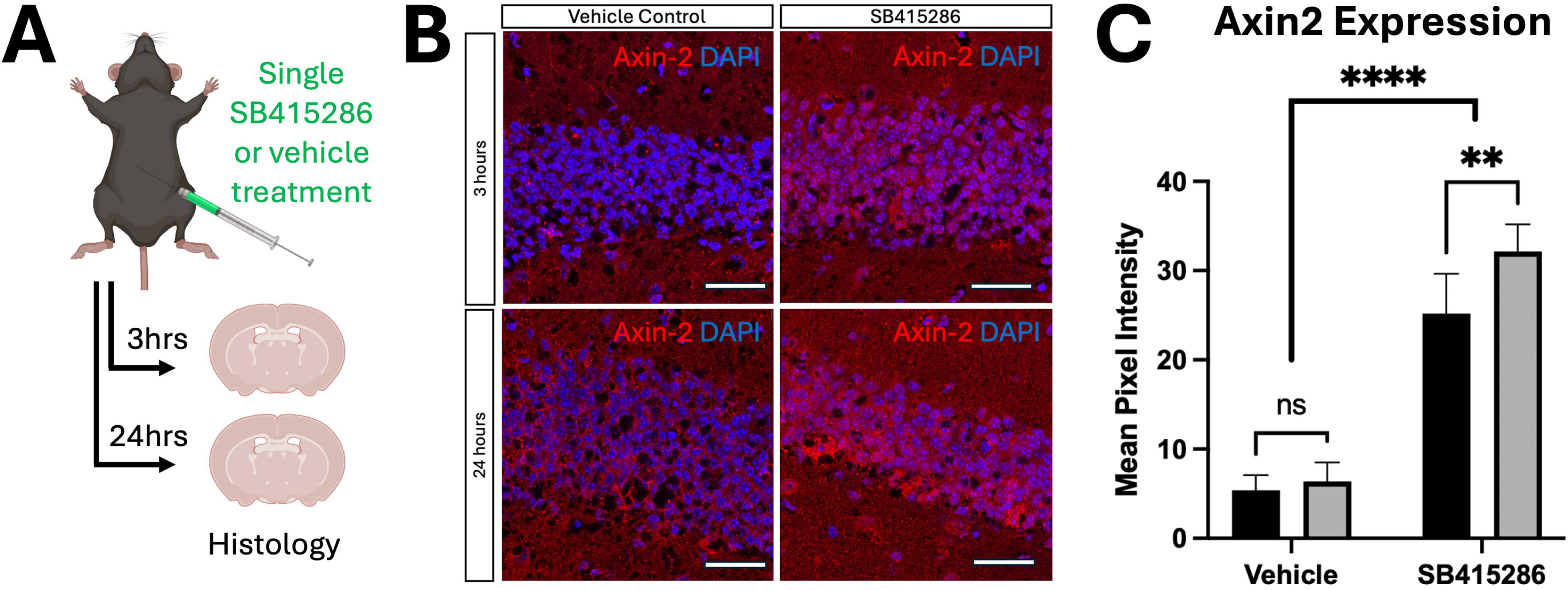
SB415286 activates Wnt signaling in dentate granule cells. A) GSK3b inhibitor SB415286 was administered to mice systemically by intraperitoneal injection, and brains were harvested 3-hours and 24-hours after a single injection. B) Wnt activity marker Axin2 (Axin2, red) was upregulated in the dentate granule cell layer (DAPI, blue) 3-hours and 24-hours after SB415286 treatment. C) Quantitative immunohistochemistry demonstrates 5±0.94-fold and 7±1.25-fold increase in Wnt activity in the innermost third of the granule cell layer 3-hours and 24-hours after SB415286 treatment compared to vehicle treated controls.

Having established that SB415286 treatment activates Wnt activity in dentate granule cells, we investigated the effect of SB415286 treatment on granule cell remodeling during epileptogenesis in two distinct models of temporal lobe epilepsy. In these experiments, mice received either intrahippocampal kainate (IHK) to induce unilateral mesial temporal lobe epilepsy with hippocampal sclerosis (Figure 2A), or intraperitoneal kainate (IPK) to induce bilateral temporal lobe epilepsy (Figure 2B). Control mice received intrahippocampal saline (IHS) or intraperitoneal saline (IPS) respectively. In each cohort, mice were treated daily for 2-weeks with either vehicle or Wnt activator SB415286, beginning 24 hours after kainate injection. This period was chosen as it corresponds to the latent period during which epileptogenesis occurs; spontaneous recurrent seizures are known to develop within 2 weeks after both intrahippocampal kainate (IHK) and intraperitoneal kainate injection (IPK) ^25,27,29,53^. Systemic treatment with Wnt activator SB415286 or vehicle began 24-hours after kainate injection to avoid potential confounding effects of drug treatment on kainate-induced status epilepticus, and to better model a translational paradigm in which a patient might receive an anti-epileptogenic treatment within 24 hours after presenting with a first seizure.

**Figure 2.**
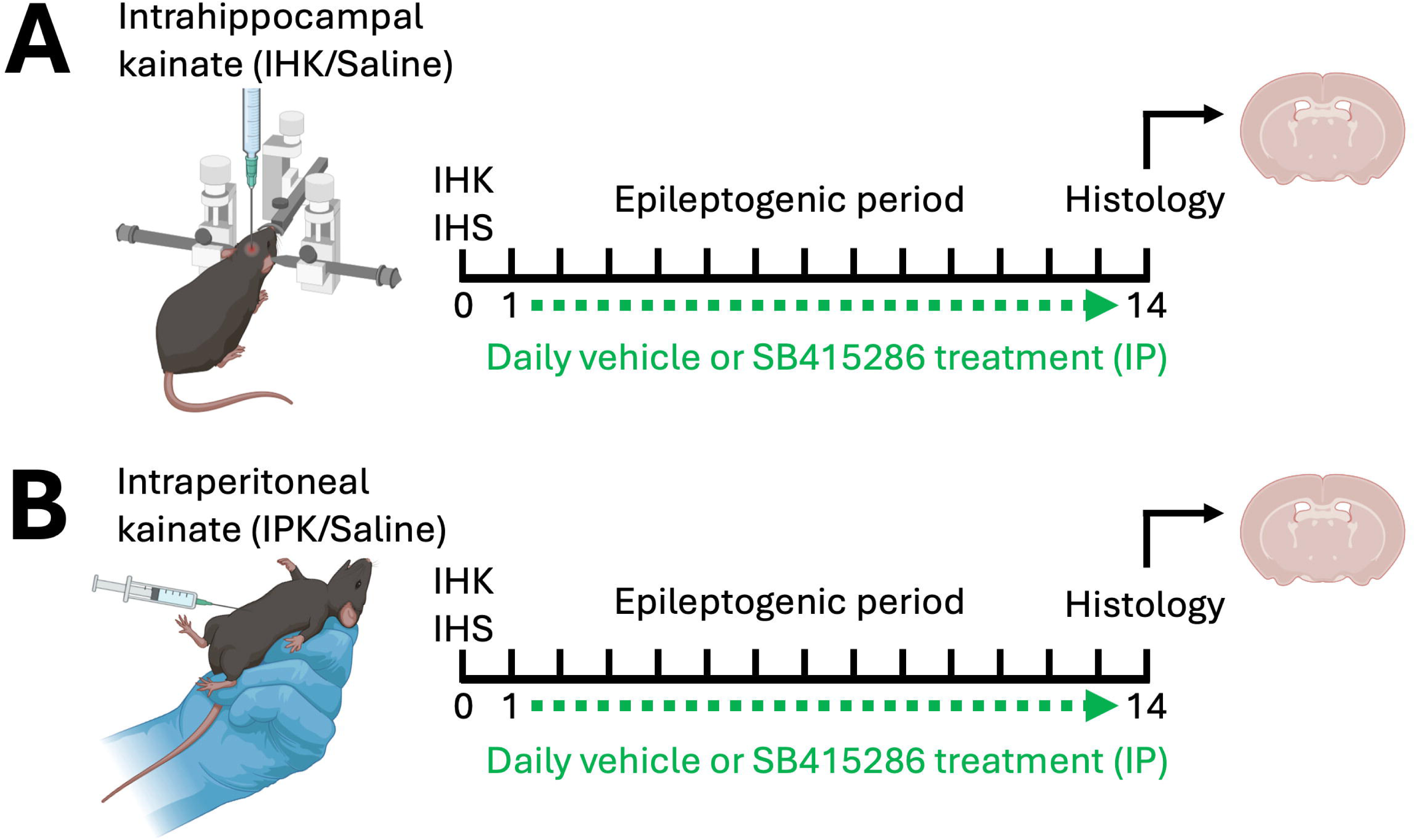
Induction of unilateral and bilateral temporal lobe epilepsy and experimental timeline. A) Unilateral temporal lobe epilepsy was induced by stereotactic injection of intrahippocampal kainate (IHK). Control mice received intrahippocampal saline (IHS). B) Bilateral temporal lobe epilepsy was induced by intraperitoneal injection of kainate (IPK). Control mice received intraperitoneal saline (IPS). For both models, mice then received daily intraperitoneal treatment with SB415286 or vehicle for 14 days, beginning 24-hours after seizure induction, prior to histological assessment.

We examined histological changes in immature dentate granule cells after IHK and IHS control in mice treated with SB415286 and vehicle for 2-weeks. Given the asymmetric nature of the unilateral intrahippocampal kainate injection mode, we examined granule cell remodeling both ipsilateral (Figure 3A) and contralateral (Figure 3B) to the site of intrahippocampal kainate injection, as it is well established that the contralateral dentate gyrus also shows changes in remodeling and excitability despite the lack of sclerosis ^27,41,54^. As we have observed previously ^27,41^, IHK resulted in the loss of immature dentate granule cell neurons and dispersion of the granule cell layer and radial migration of the remaining granule cells in the ipsilateral dentate gyrus of the hippocampus in vehicle treated animals compared to intrahippocampal saline (IHS) control mice (Figure 4 p<0.0001). When animals were treated with Wnt activator SB415286, granule cell dispersion (Figure 4A, p<0.01) and granule cell radial migration (Figure 4b, p<0.001) were significantly reduced in the ipsilateral dentate gyrus after IHK; there were no changes in granule cell morphology in IHS control mice treated with Wnt activator SB415286 compared to vehicle (Figure 3A). As described previously ^27,41^, we again observed immature granule cell proliferation in the contralateral dentate gyrus after IHK (Fig 4C, p<0.0001); this was also prevented by Wnt activator SB415286 treatment (Figure 4C, p<0.05). There were no changes in immature dentate granule cell dispersion or migration after IHK in the contralateral dentate gyrus (Figure 4A, 4B). Change in proximal dendrite angle of immature dentate granule cells is also an established marker of epileptogenesis and has been used as a marker of anti-epileptogenic treatment response in rodent models ^27,43^. We therefore also examined the proximal dendrite angle of POMC-GFP^+^ immature dentate granule cells after IHK in mice treated with Wnt activator SB415286 and vehicle control (Figure 4D, 4E). We observed an increase in proximal dendrite angulation towards the hilus after IHK in both the ipsilateral (p<0.001) and contralateral dentate gyrus (p<0.001) compared to intrahippocampal saline injected controls. When kainate injected epileptic animals were treated with Wnt activator SB415286, pathological proximal dendrite angulation was prevented (p<0.01) with the distribution approximating intrahippocampal injected saline controls. In intrahippocampal saline injected animals, SB415286 treatment had no effect on proximal dendrite angle in the ipsilateral dentate gyrus; proximal dendrite angle was reduced in the contralateral dentate gyrus by SB415285 treatment compared to vehicle control treatment. Example dendrite tracings in the IHK/IHS experimental subgroups are provided in Supplemental Figure 1.

**Figure 3.**
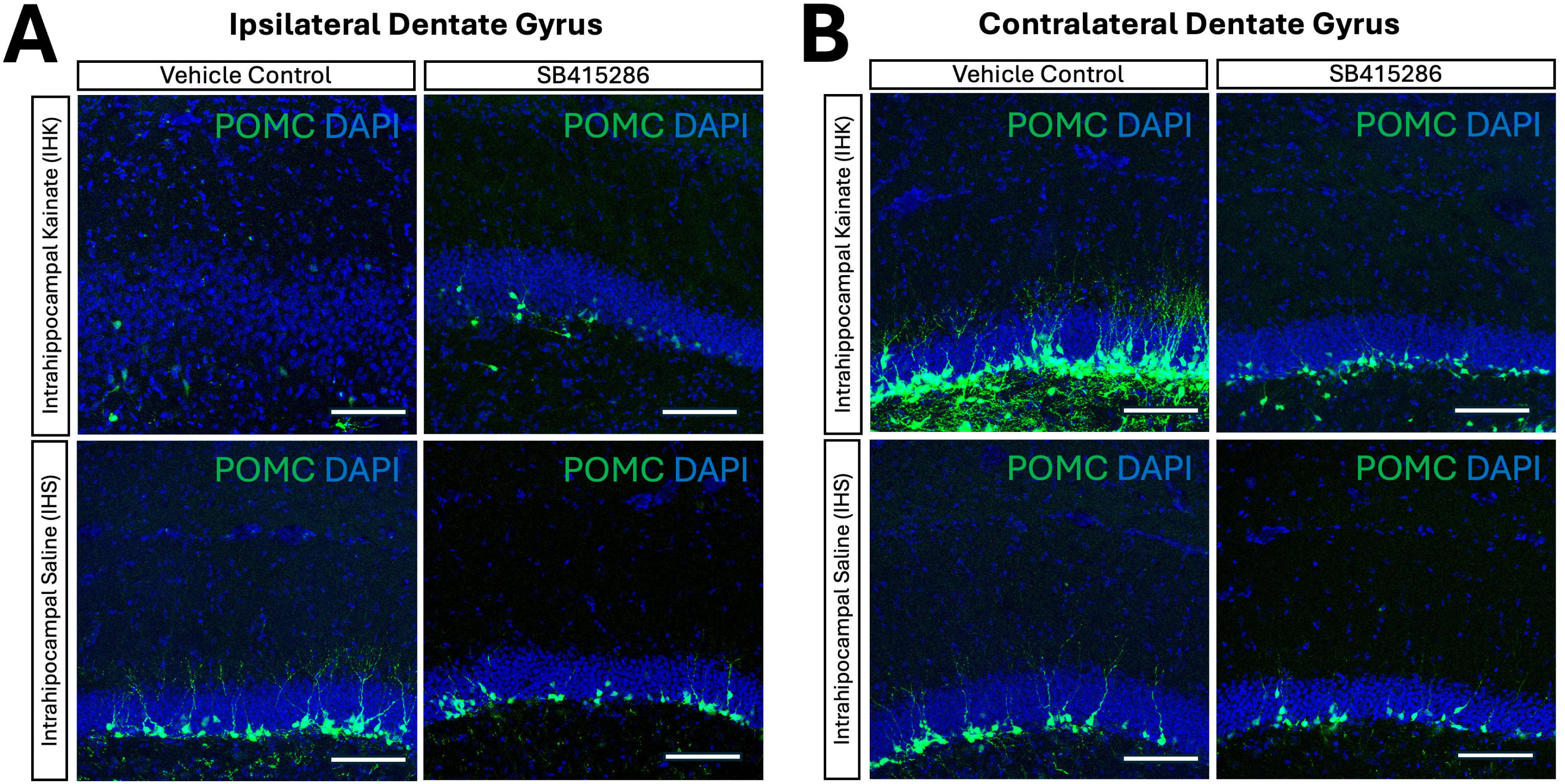
Immature dentate granule cell remodeling in the intrahippocampal kainate (IHK) model of unilateral TLE. A) Immunohistochemistry of the dentate gyrus ipsilateral to IHK injection demonstrates loss of immature dentate granule cells (POMC-GFP, green) and granule cell dispersion (DAPI, blue) 2-weeks after seizure induction by IHK in vehicle treated animals compared to intrahippocampal saline controls. SB415286 treatment after IHK rescued granule cell dispersion and immature dentate granule cell loss. B) Immunohistochemistry of the dentate gyrus contralateral to IHK injection demonstrates aberrant neurogenesis and dendritic arbor formation after IHK in vehicle treated animals compared to intrahippocampal saline controls. SB415286 treatment rescued the aberrant neurogenesis and dendritic arbor outgrowth observed after IHK. Scale bar 100µm.

**Figure 4.**
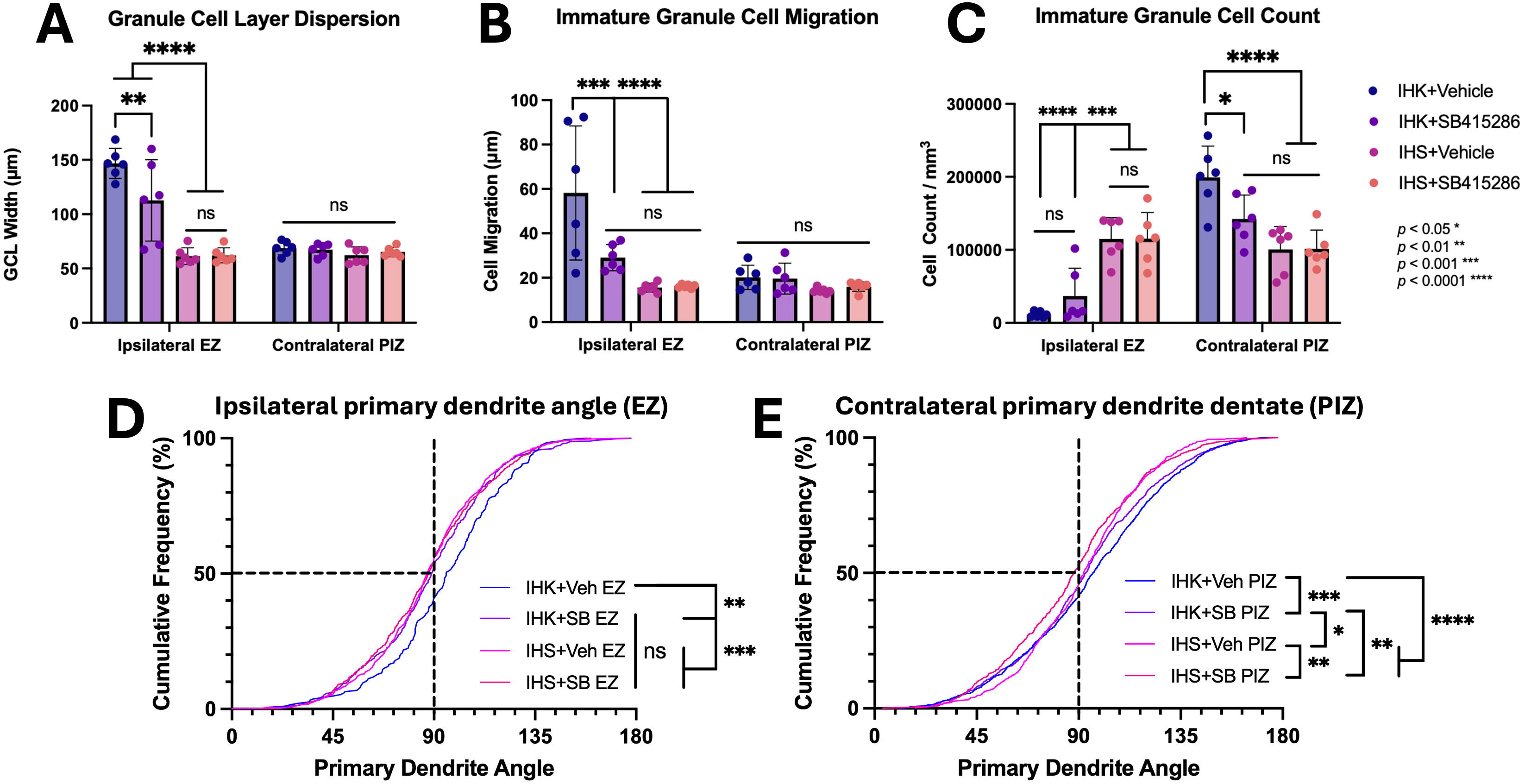
Quantification of immature dentate granule cell epileptogenic remodeling in the IHK model of unilateral temporal lobe epilepsy. A) Granule cell layer dispersion increased in the ipsilateral dentate gyrus after IHK compared to saline controls. SB415286 treatment significantly reduced granule cell dispersion after IHK compared to vehicle treated mice. There were no significant changes in granule cell dispersion in the contralateral dentate gyrus after IHK or with SB415286 treatment. B) Immature dentate granule cell migration increased in the ipsilateral dentate gyrus after IHK compared to saline controls. SB415286 also significantly reduced granule cell migration after IHK compared to vehicle treated mice, to levels comparable to saline controls. There were also no significant changes in granule cell migration in the contralateral dentate gyrus. C) IHK caused significant granule cell loss in the ipsilateral dentate gyrus compared to saline controls; this was unaffected by SB415286 treatment. In the contralateral dentate gyrus, IHK increased neurogenesis compared to saline controls. SB415286 treatment significantly reduced aberrant neurogenesis after IHK compared to vehicle treated mice. Granule cell proximal dendrite angle increased after IHK in both the D) ipsilateral and E) contralateral dentate gyrus. In both regions, SB415286 treatment reduced proximal dendrite angle orientation to levels comparable with saline control mice.

Bilateral TLE is a distinct clinical entity from unilateral TLE, with differing neurosurgical treatment paradigms, and merits independent evaluation from unilateral TLE. The intraperitoneal kainate model (IPK) uses systemically administered kainate to induce TLE, rather than focal intrahippocampal administration in the IHK model. As kainate is applied systemically, both dentate gyri are affected equally, resulting in a TLE model that more closely approximates bilateral hippocampal TLE. We examined immature dentate granule cell remodeling after IPK model and intraperitoneal saline (IPS) control in POMC-GFP^+^ mice, treated with vehicle and Wnt activator SB415286 daily for 2-weeks (Fig 5). In contrast to the IHK model, the granule cell layer did not undergo dispersion after IPK (Fig 6A). Immature granule cell migration increased after IPK (*p*<0.01, Fig 6B), which was also observed in the ipsilateral dentate gyrus after IHK. Immature dentate granule cell neurogenesis also increased after IPK (*p*<0.01, Fig 6C), which was also observed in the contralateral dentate gyrus after IHK. Wnt activation with SB416286 had no effect on granule cell layer dispersion, granule cell migration and granule cell count after IPK or IPS (Fig 6A-C). We also examined proximal dendrite angle after IPK. Similar to IHK, we observed an increase in proximal dendrite angle of immature dentate granule cells after IPK in vehicle treated animals. We also observed that pro-epileptogenic remodeling of proximal dendrites was prevented with Wnt activation after IPK (Fig 6D). Example dendrite tracings in the IPK/IPS experimental subgroups are provided in Supplemental Figure 2. This finding further suggests that dentate granule cell proximal dendrite angle is a valid and reliable marker of epileptogenesis in multiple temporal lobe epilepsy models.

**Figure 5.**
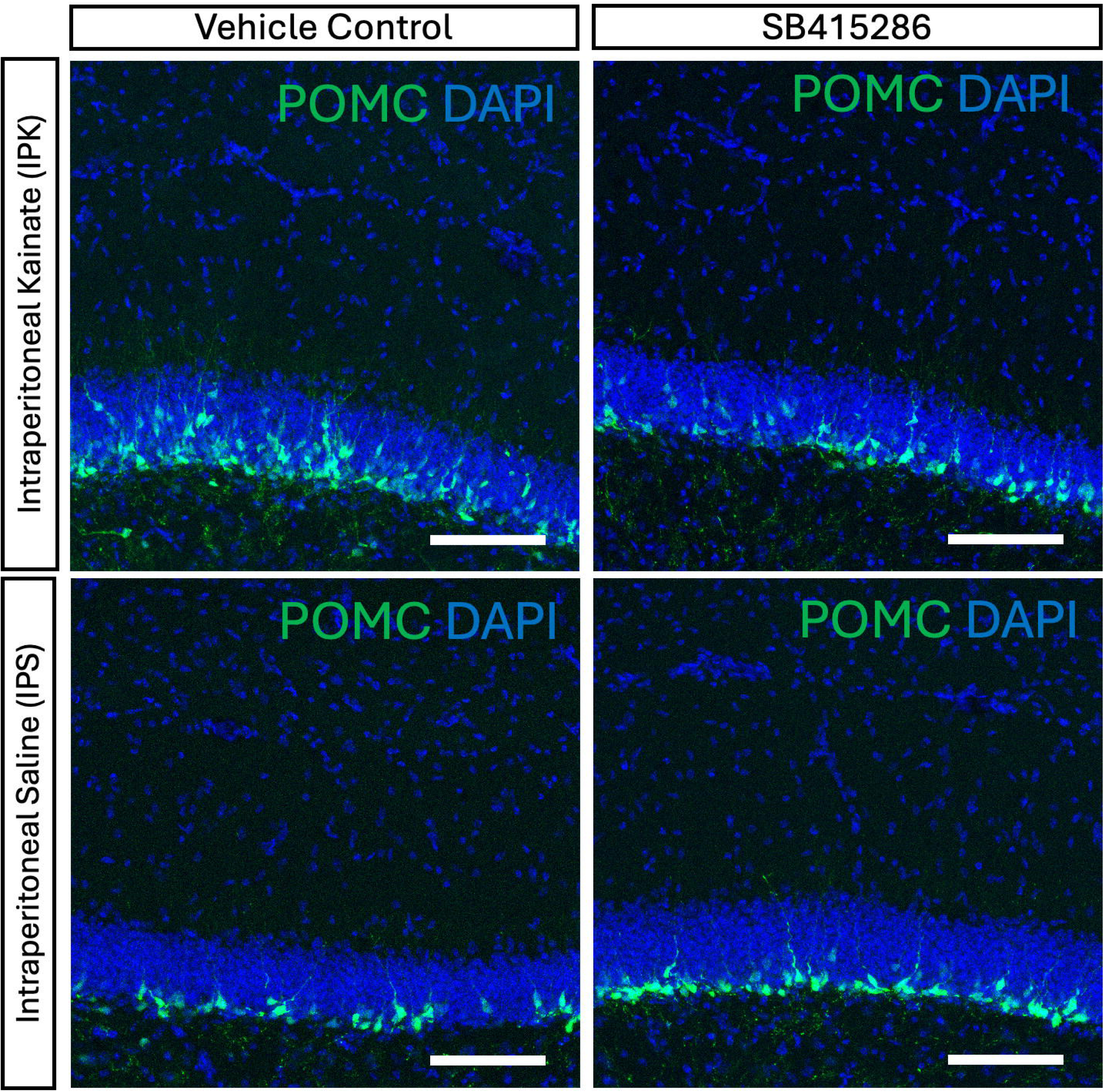
Immature dentate granule cell remodeling in the intraperitoneal kainate (IPK) model of bilateral TLE. Immunohistochemistry of the dentate gyrus demonstrates increased neurogenesis and migration of immature dentate granule cells (POMC-GFP, green) 2-weeks after seizure induction by IPK compared to intraperitoneal saline (IPS) injected control animals; granule cell layer dispersion is unchanged in epileptic mice in this model. SB415286 treatment did not alter granule cell layer dispersion, neurogenesis or immature dentate granule cell migration in IPK injected epileptic animals or IPS controls. The right dentate gyrus is demonstrated; there were no differences between the left and right hemisphere after IPS/IPK. Scale bar 100µm.

**Figure 6.**
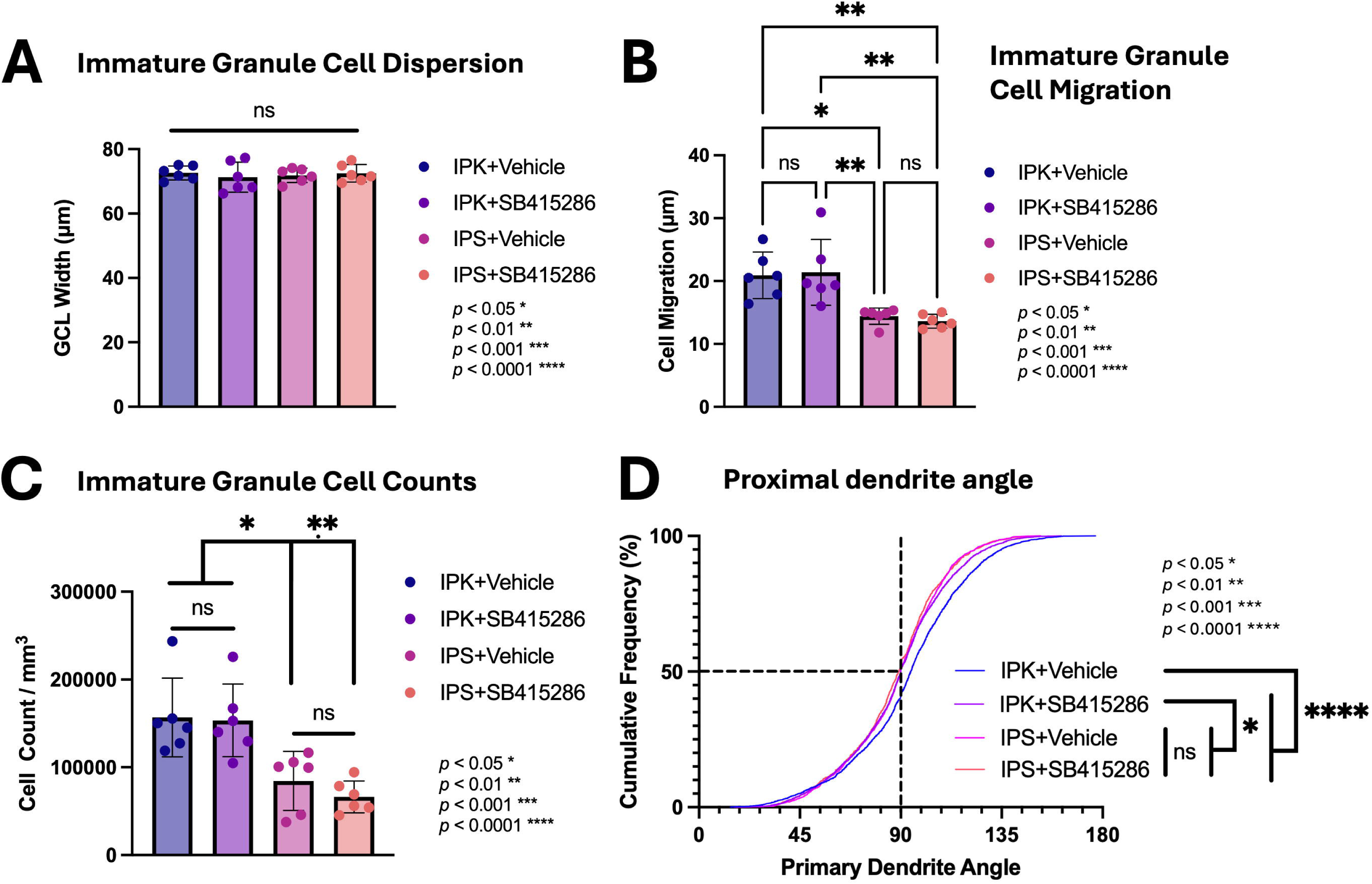
Quantification of immature dentate granule cell epileptogenic remodeling in the IPK model of bilateral temporal lobe epilepsy. A) Granule cell dispersion was unchanged 2-weeks after seizure induction with IPK compared to IPS controls and was unaffected by SB415286 treatment in both epileptic and control animal groups. B) Immature dentate granule cell migration increased in epileptic animals 2-weeks after IPK compared to IPS injected control animals. SB415286 treatment did not alter immature dentate granule cell migration in epileptic or control animals. C) Dentate granule cell neurogenesis increased in epileptic animals 2-weeks after IPK compared to IPS injected control animals. SB415286 treatment did not alter immature dentate granule cell neurogenesis in epileptic or control animals. D) Granule cell proximal dendrite angle increased after IPK. SB415286 treatment reduced proximal dendrite angle orientation to levels comparable with saline control mice.

In the following experiments, we sought to examine Wnt activity in both models of TLE and in response to Wnt activator SB415286 treatment by quantifying Axin2 expression in POMC^+^ adult-born dentate granule cells, to better understand the mechanistic role of Wnt activation in granule cell remodeling. In the IHK model of unilateral TLE, in the ipsilateral dentate gyrus, Wnt signaling activity reduced significantly below baseline levels 2-weeks after IHK in vehicle treated animals. When IHK-injected mice were treated with Wnt activator SB415286, Wnt signaling levels increased to baseline levels observed in saline control animals (Figure 7A-B). In the hemisphere contralateral to IHK injection, Wnt signaling levels in immature dentate granule cells were unchanged after IHK, compared to intrahippocampal saline controls (Figure 7C-D). As expected, SB415286 treatment in intrahippocampal saline injected control mice increased Wnt activity above baseline in both hemispheres. In the IPK model of bilateral TLE, Wnt signaling activity also decreased below baseline in dentate granule cells in untreated epileptic mice and SB415286 administration increased Wnt activity in these cells to baseline levels observed in saline controls (Figure 8A-B). As expected, SB415286 treatment in saline control mice increased Wnt activity above baseline. Notably, both dentate gyri in the IPK model of bilateral TLE showed similar Wnt signaling changes (Supplemental Figure 3), whereas Wnt signaling changes differed between hemispheres in the IHK model of unilateral TLE. Notably, Wnt signaling changes were similar between the bilateral dentate gyri after IPK and the ipsilateral dentate gyrus after IHK, representing the seizure onset zones in the bilateral and unilateral TLE respectively. These data highlight the differences between the two models and further suggest that Wnt signaling changes occur during epileptogenesis specific to the epileptogenic zone.

**Figure 7.**
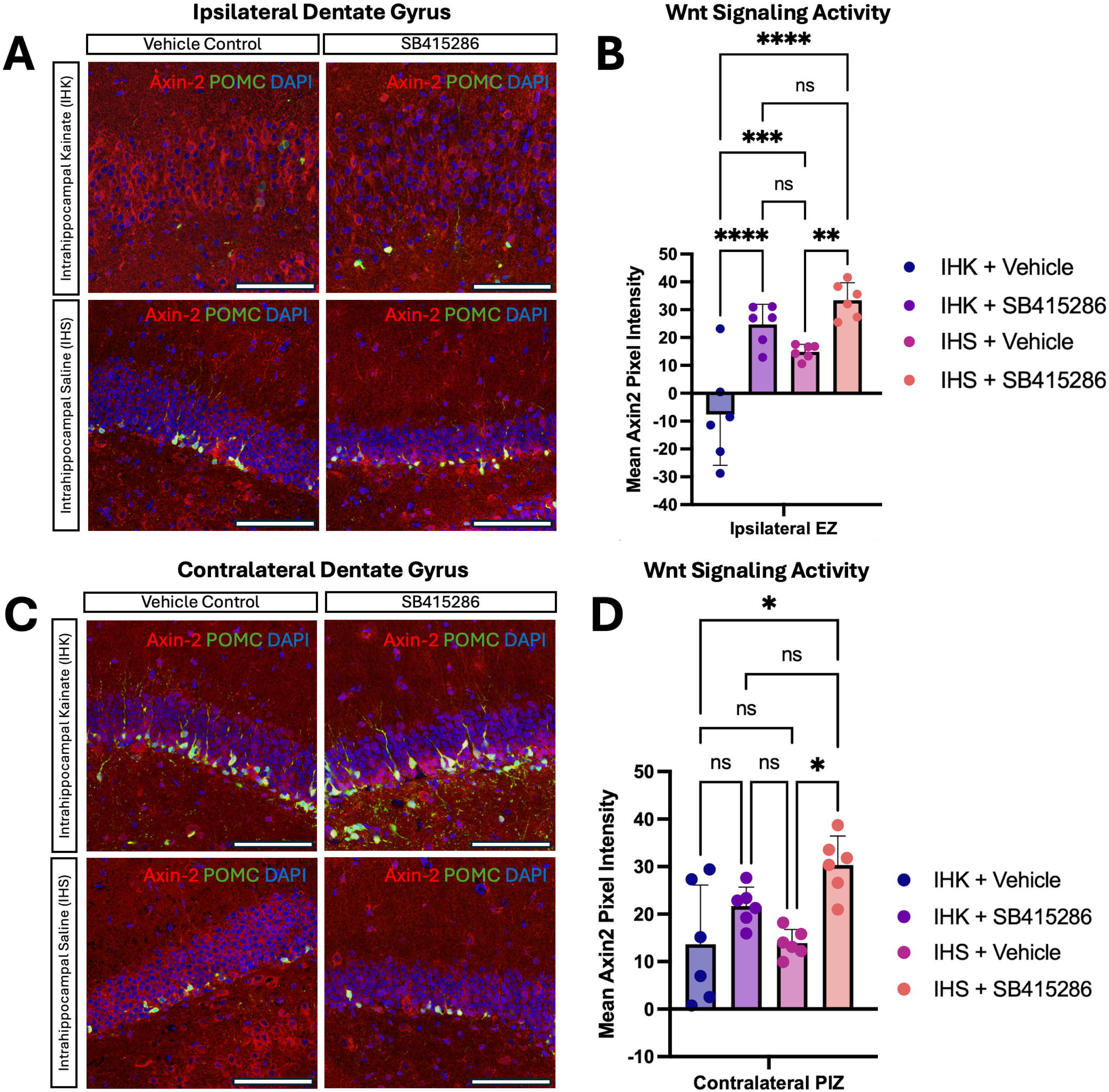
Wnt activator SB415286 increases Wnt signaling in immature dentate granule cells in the ipsilateral dentate gyrus after IHK. Wnt activator SB415286 increases Wnt signaling in immature dentate granule cells after IHK. Wnt signaling levels in POMC-GFP^+^ immature dentate granule cells (POMC-GFP, green) were determined by Axin2 co-immunostaining (Axin2, red) in mice with unilateral temporal lobe epilepsy after IHK and IHS injected control animals after vehicle and SB415286 treatment. A) Immunofluorescence images for the ipsilateral dentate gyrus, scale bar 100µm. B) Quantification of Axin2 fluorescence intensity in POMC-GFP^+^ co-labelled immature dentate granule cells in the ipsilateral dentate gyrus demonstrated that Wnt signaling levels decreased below baseline in immature dentate granule cells after IHK. SB415286 treatment increased Wnt signaling levels in immature dentate granule cells after IHK to baseline IHS control levels. In IHS control animals, SB415286 increased Wnt signaling in immature dentate granule cells as expected. C) Immunofluorescence images for the contralateral dentate gyrus, scale bar 100µm. D) In the contralateral dentate gyrus, SB41586 increased Wnt signaling levels in IHS injected control mice as expected; there were no changes in Wnt signaling levels after IHK.

**Figure 8.**
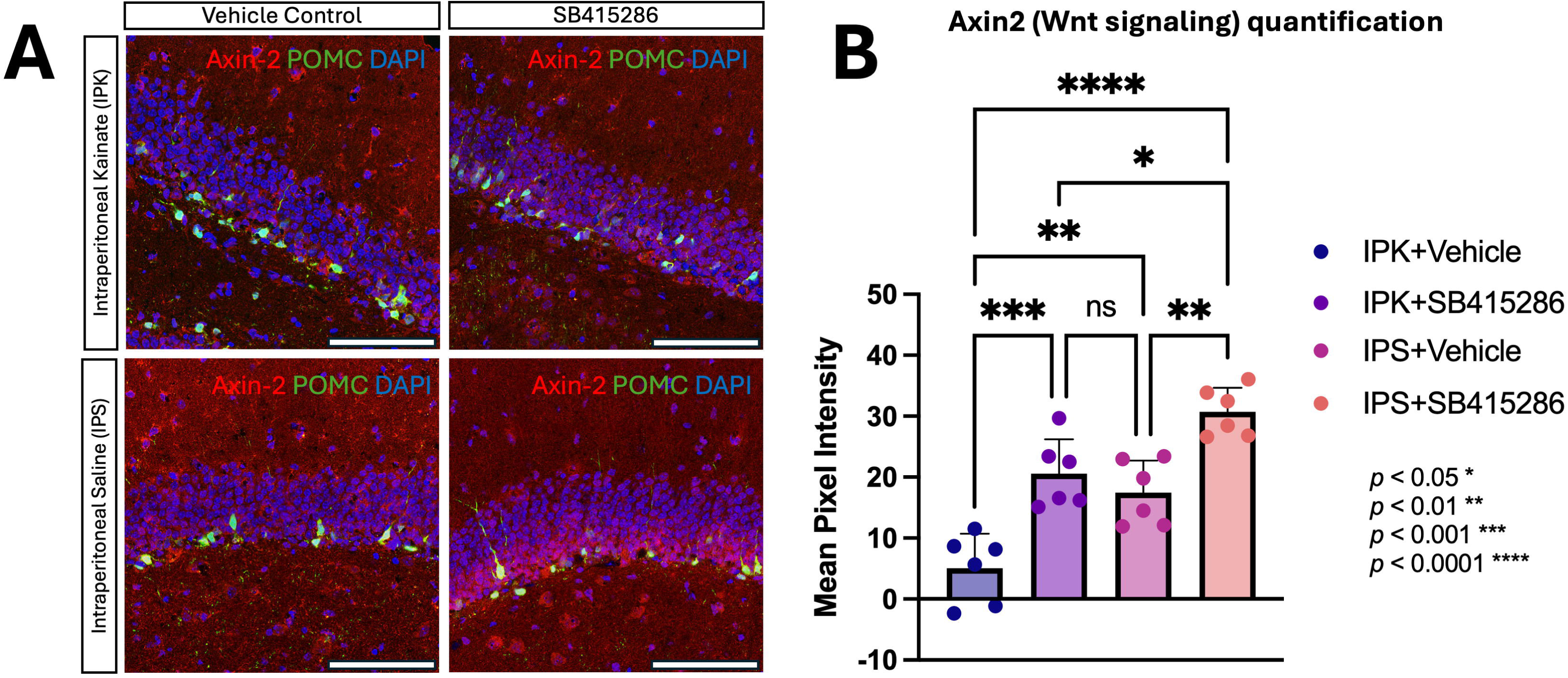
Wnt activator SB415286 increases Wnt signaling in immature dentate granule cells after IPK. A) Wnt signaling in POMC-GFP^+^ immature dentate granule cells (POMC-GFP, green) was determined by Axin2 co-immunostaining (Axin2, red) in mice with bilateral temporal lobe epilepsy after IPK and IPS injected control animals after vehicle and SB415286 treatment, scale bar 100µm. B) Quantification of Axin2 fluorescence intensity in POMC-GFP^+^ co-labelled immature dentate granule cells demonstrated that Wnt signaling levels decreased below baseline in immature dentate granule cells after IPK. SB415286 treatment after IPK increased Wnt signaling levels in immature dentate granule cells to IPS-injected control levels. In IPS injected control animals, SB415286 increased Wnt signaling in immature dentate granule cells as expected.

These findings demonstrate that Wnt signaling decreases in adult-born granule cells during epileptogenesis, and that increased Wnt activity after SB415286 in remodeled dentate granule cells is associated with prevention of epileptogenic remodeling in both unilateral and bilateral temporal lobe epilepsy models. These data therefore suggest that Wnt activation could be developed as a novel therapeutic approach for both forms of clinical epilepsy.

## Discussion

In this study, we evaluated the role of Wnt signaling in epileptogenic remodeling of immature dentate granule cells using two models of temporal lobe epilepsy, the intrahippocampal kainate model of unilateral temporal lobe epilepsy (IHK) and the intraperitoneal kainate model of bilateral temporal lobe. We demonstrate that both the IHK and IPK models bear histological hallmarks of epileptogenesis, and that canonical Wnt signaling is reduced in immature dentate granule cells during epileptogenesis. We further demonstrate that systemic treatment with canonical Wnt activator SB415286 prevents pathological immature dentate granule cell remodeling in both models and restores Wnt signaling in these cells to baseline levels. These experiments therefore suggest that Wnt signaling plays a role in granule cell remodeling during epileptogenesis in temporal lobe epilepsy, and that Wnt activation has translational potential as an anti-epileptogenic therapy.

The IHK model of unilateral TLE demonstrated well-described histological derangements of immature dentate granule cell morphology. As other studies have observed previously, IHK results in extensive granule cell remodeling, including cell loss and migration, granule cell layer dispersion, mossy fiber sprouting and proximal dendrite angulation in the ipsilateral hemisphere ^37–39,55,56^. Uniquely, in our previous work we also investigated the contralateral hemisphere and discovered that this hemisphere also undergoes epileptogenic remodeling, including changes in dendritic arborization, proximal dendrite angle and granule cell neurogenesis after IHK ^27,41^. These findings are accurately reproduced in this study, demonstrating the reliability and reproducibility of this model for investigating unilateral TLE and testing novel candidate therapies. In the IPK model, early studies using single intraperitoneal kainate doses reported variable effectiveness, with single doses up to 35mg/kg resulting in either high mortality or lack of neurodegeneration ^22–24^, though some reports describe histological changes after single dose IPK ^57,58^. With newer repeated low-dose protocols, as used in our study, IPK robustly induces status epilepticus with long-term electrographic seizures up to 18 weeks after IPK ^25,26^. However, histological changes and granule cell remodeling in this model have not previously been described. In this study, we provide a novel description of histological markers of epileptogenesis in the repeated low dose intraperitoneal kainate model of bilateral TLE; these include pathological alterations in immature dentate granule cell neurogenesis, migration and proximal dendrite angle after IPK, without granule cell layer dispersion, symmetrically in both dentate gyri. Notably, the IPK model does not directly reproduce the histological characteristics of the ipsilateral dentate gyrus after IHK, highlighting that these models are indeed distinct entities. There is, however, some overlap between these models, with the IPK model sharing aspects of both the ipsilateral dentate gyrus after IHK, such as granule cell migration and alteration of proximal dendrite angle, and the contralateral dentate gyrus after IHK, such as neurogenesis and the lack of granule cell layer dispersion. These findings suggest that pathological remodeling of immature dentate granule cells may contribute to epileptogenesis in both forms of temporal lobe epilepsy and that underlying mechanisms may be shared, thus representing a common therapeutic target for both forms of TLE.

We investigated the role of Wnt signaling in the development of unilateral and bilateral TLE, examining both changes in hippocampal Wnt signaling during epileptogenesis and the effect of therapeutic Wnt modulation on epileptogenic remodeling. We observed that in both the IHK and IPK models, canonical Wnt signaling marker Axin2 expression decreased in immature dentate granule cells. In the IPK model, this was consistent across both hemispheres, in the IHK model this reduction in Wnt signaling in epileptic animals was only observed in the ipsilateral dentate gyrus. This finding is also consistent with our previous work that independently demonstrated that Wnt signaling reduces in the hippocampal dentate gyrus during epileptogenesis ^32^. These findings suggest that Wnt signaling decreases specifically in the epileptogenic zone during epileptogenesis and therefore may play a critical role in the etiopathogenesis of TLE. The finding that Wnt signaling decreases specifically in dentate granule cells in the epileptogenic zone led us to hypothesize that restoring Wnt signaling may have an anti-epileptogenic effect. We sought to examine this hypothesis by testing Wnt activator SB415286 as a novel anti-epileptogenic therapy for TLE. Mice were treated with SB415286 for the duration of the epileptogenic period, with treatment beginning 24-hours after seizure induction. This delay in treatment initiation was chosen to more accurately test a clinical paradigm whereby patients are treated within 24-hours of first seizure. Wnt activator SB415286 treatment prevented pro-epileptogenic granule cell remodeling in both the IHK and IPK models of TLE. In the IHK model, SB415286 treatment reduced granule cell layer dispersion and granule cell migration in the ipsilateral dentate gyrus, and aberrant neurogenesis in the contralateral dentate gyrus. Wnt activator treatment also prevented remodeling of the proximal dendrite of immature dentate granule cells, a key epileptogenic change observed in other TLE models ^43^. In the IPK model, SB415286 treatment also prevented remodeling of the proximal dendrite in both hemispheres, with findings comparable to the ipsilateral dentate gyrus after IHK. These findings exceeded those observed in our previous work testing Wnt activator Chir99021 after IHK ^27^; this difference may reflect the longer half-life of SB415286 ^47^, which, with reduced off-target effects ^36^, may translate to greater clinical translational potential.

We also quantified changes in Wnt signaling activity in immature dentate granules with SB415286 treatment. We observed that SB415286 treatment increased Wnt signaling activity in dentate granule cells in control animals above baseline levels. In epileptic animals, however, Wnt signaling activity in dentate granule cells was restored to baseline in both IHK and IPK models and did not rise above physiological levels. These findings suggest that Wnt activators may be applied safely during epileptogenesis. Notably, in our prior work we observed that saline control mice treated with Wnt activator Chir99021 demonstrated impaired cognitive testing compared to vehicle treated controls, as opposed to epileptic mice which improved with treatment; this finding may reflect a supra-physiological increase in hippocampal Wnt signaling in non-epileptic control mice, though this remains to be determined in future experiments. This suggests that timing of Wnt activator treatment may also be critical for clinical translation. Furthermore, changes in Wnt signaling were similar in both dentate gyri in bilateral TLE and similar to the epileptogenic zone in unilateral TLE, suggesting that Wnt signaling changes we observed are specific to the epileptogenic zone, further highlighting a shared therapeutic mechanism for Wnt signaling augmentation across both models of TLE. This is also valuable for clinical translation as, since risk factors for unilateral and bilateral TLE are shared, the same anti-epileptogenic intervention could be effective against both forms of TLE.

In conclusion, these data strongly support an etiological model in which an epileptogenic event results in the loss of Wnt signaling in adult-born immature dentate granule cells in the epileptogenic zone, resulting in pathological remodeling of these cells and the development of epileptic circuits in temporal lobe epilepsy. Our data further suggest that restoring Wnt signaling in this critical time-period could be a novel therapeutic strategy for preventing epileptogenesis and temporal lobe epilepsy. These findings therefore have significant translational implications. Importantly, our investigation of two independent models of TLE allows comparisons to be drawn. Unilateral and bilateral hippocampal TLE are common forms of clinical temporal lobe epilepsy and share common risk factors ^16–20^. As a result, in clinical epilepsy management, it may be challenging to accurately determine which form of TLE patients might develop after an epileptogenic exposure. Anti-epileptogenic interventions for MTLE may therefore need provide preventative efficacy for both unilateral and bilateral forms of TLE, as risk factors alone may be insufficient to determine which form of MTLE may ensue. In this study, we demonstrate that Wnt activation effectively reduces epileptogenic remodeling in both models of TLE, suggesting that Wnt activation may represent a novel therapeutic candidate for the prevention of both unilateral and bilateral TLE in patients at high-risk of epileptogenesis. Notably, Wnt activator Chir99021 is currently in human clinical trials for sensorineural hearing loss ^59^, highlighting the therapeutic potential of Wnt activating compounds. In future studies, it will be critical to determine whether the prevention of pathological remodeling is also reflected in a durable reduction in electrographic and clinical epileptic activity. Safety data will also be important for clinical translation and pilot clinical studies.

Supplemental Figure 1. Representative dendrite traces of immature dentate granule cells after IHK and IHS in unilateral TLE. In this panel, representative dendrite traces are provided for POMC-GFP^+^ immature dentate granule cells from the ipsilateral and contralateral hemisphere for IHK epileptic animals and IHS control animals treated with vehicle and SB415286. These tracings accompany data in figures 4D and 4E.

Supplemental Figure 2. Representative dendrite traces of immature dentate granule cells after IPK and IPS in bilateral TLE. In this panel, representative dendrite traces are provided for POMC-GFP+ immature dentate granule cells for IPK epileptic animals and IPS control animals treated with vehicle and SB415286. These tracings accompany data in figures 6D.

Supplemental Figure 3. Wnt signaling levels in adult-born dentate granule cells change symmetrically in both hemispheres in the IPK model of bilateral TLE. A) Quantification of Wnt activity marker Axin2 in POMC-GFP^+^ immature dentate granule cells demonstrated no significant differences between the left and right hemispheres in the IPK model of bilateral TLE. B) There were also no significant differences in the number of Axin2 / POMC-GFP^+^ co-positive cells in each experimental condition in the IPK/IPS model of bilateral TLE.

## Acknowledgements

This work was supported by funding from the National Institute of Neurological Disorders and Stroke (NINDS) at the National Institutes of Health (NIH) (K12NS080223), the Medical College of Wisconsin Research Affairs Committee and the Medical College of Wisconsin Department of Neurosurgery.

## Author contributions

Conceptualization, KG; Investigation, CH, NR, KG; Supervision, KG; Writing – original draft, KG; Funding acquisition, KG; Writing – review and editing, CH, NR, KG.

## Resource Availability

Lead Contact: Requests for further information and resources should be directed and will be fulfilled by the lead contact, Dr Kunal Gupta MD PhD. Materials availability: This study did not generate new unique reagents. Data and code availability: 1 Data: All data reported in this paper will be shared by the lead contact upon request. 2 Code: This paper does not report original code. 3 Additional information: Any additional information required to reanalyze the data deported in this paper is available from the lead contact upon request.

