## Supplementary figures and images for "Wnt activation prevents epileptogenic hippocampal remodeling in animal models of unilateral and bilateral temporal lobe epilepsy"

### Supplementary Figure 1

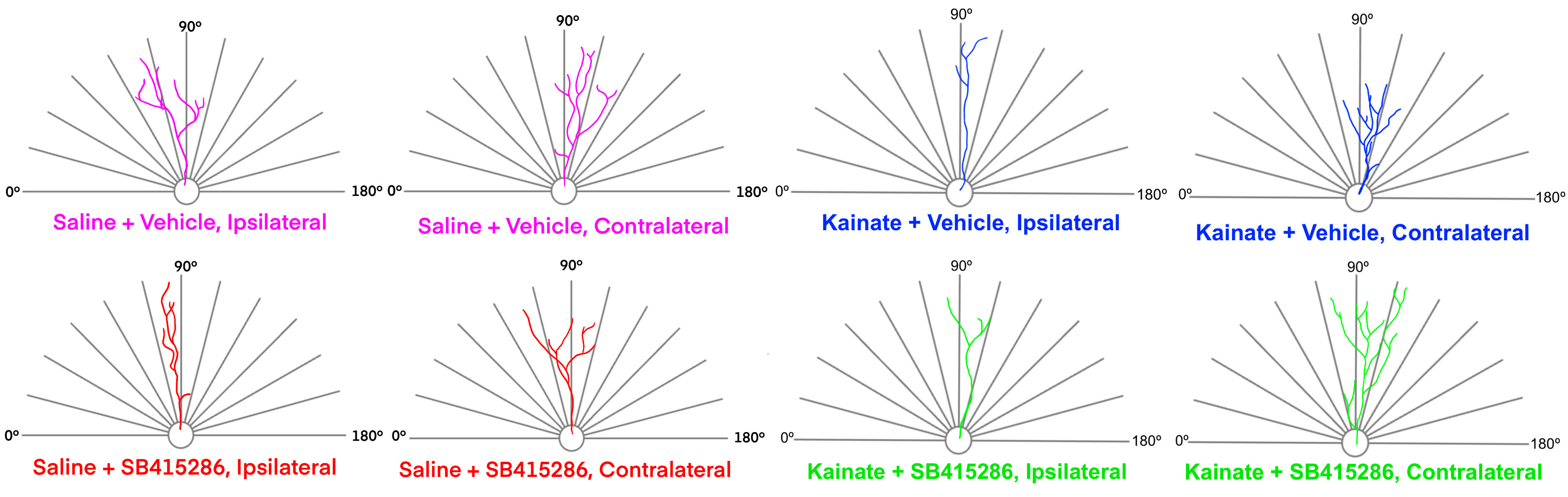

### Supplementary Figure 2

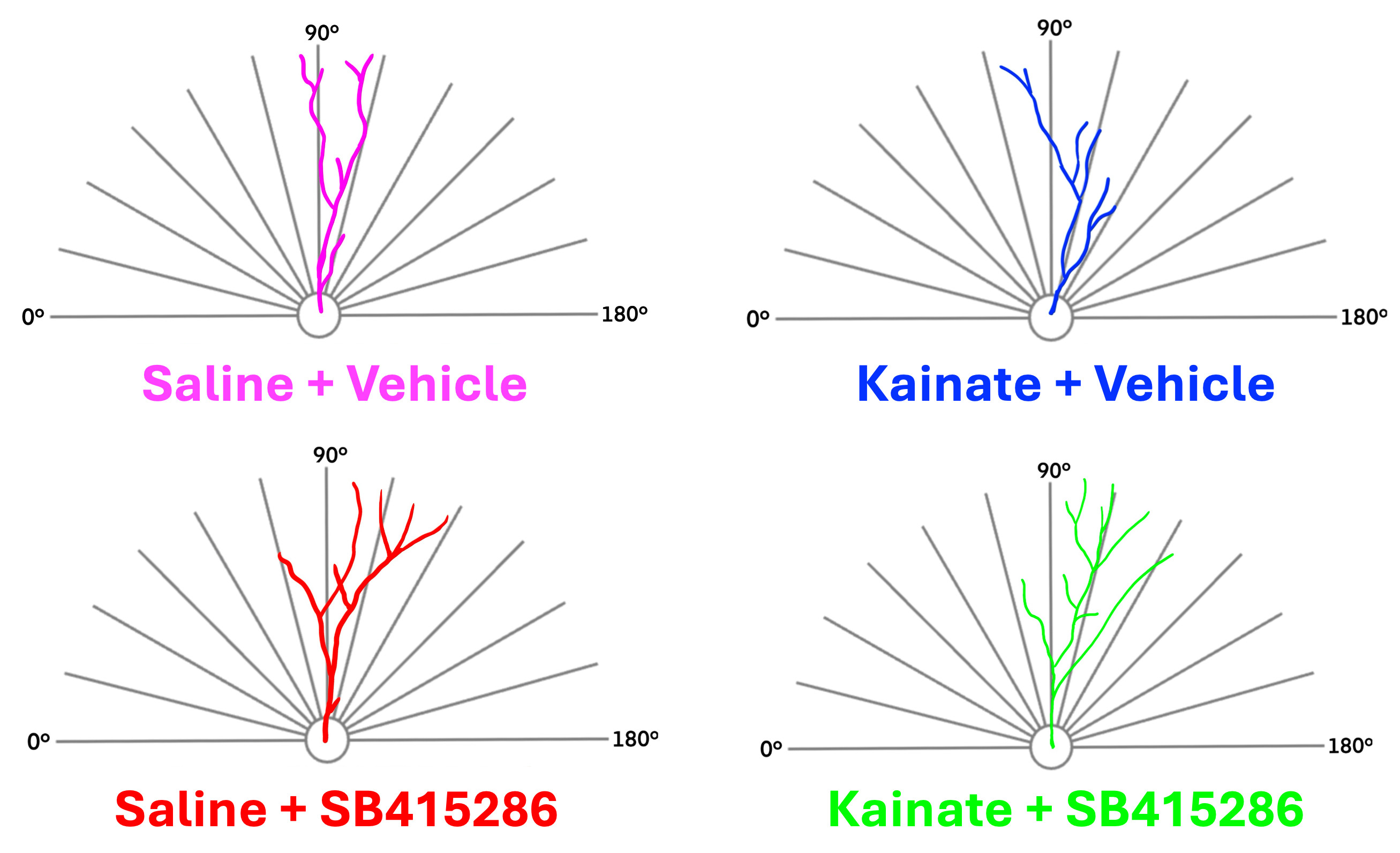

### Supplementary Figure 3

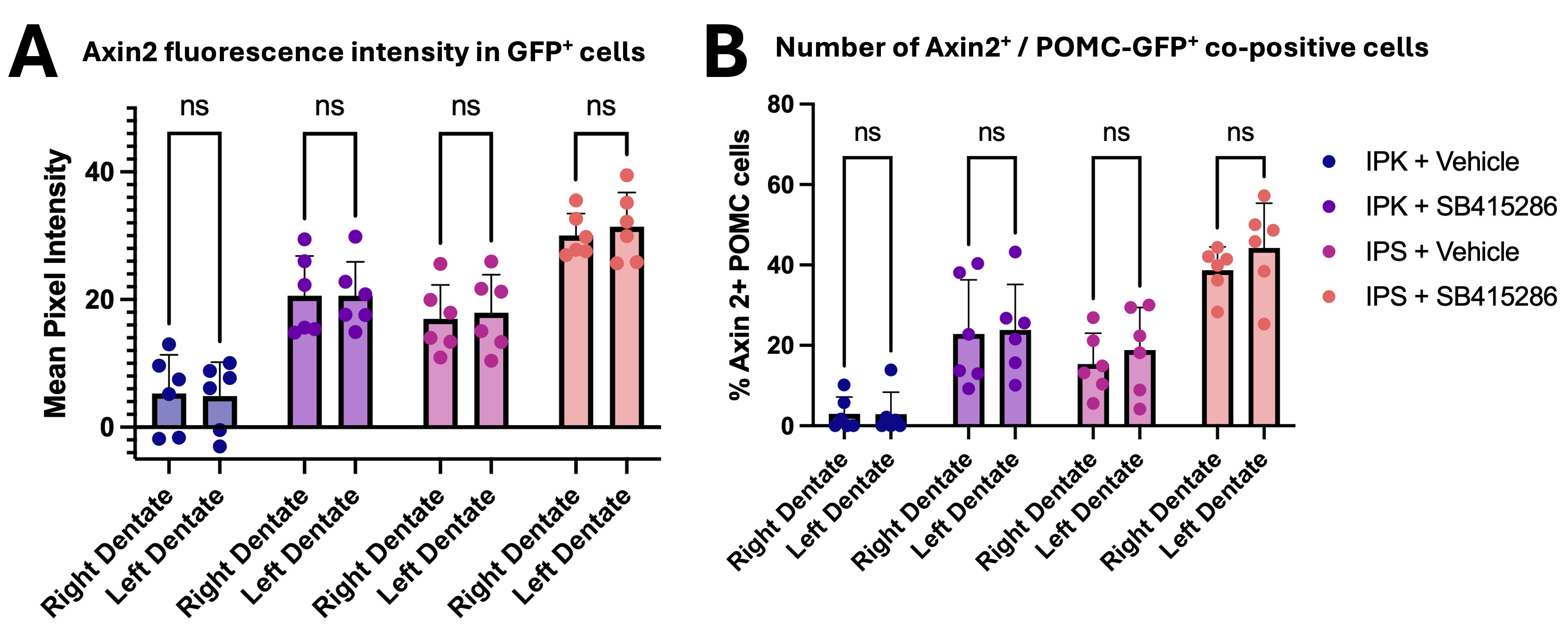
